# Structure-Guided Design and Dynamic Evaluation of VP4-Targeting siRNAs Against Rotavirus A

**DOI:** 10.64898/2026.04.03.716385

**Authors:** Abu Nayeem Ahmed, Khaleda Jahan Satu, A.B.Z. Naimur Rahman, SM Sajid Hasan, Mohammad Nazmus Sakib, Md. Ekram Hossan, Arittra Bhattacharjee, Zeshan Mahmud Chowdhury, Ziaul Faruque Joy, Md. Jahirul Islam, Mohammad Uzzal Hossain

**Affiliations:** Department of Genetic Engineering and Biotechnology, Shahjalal University of Science and Technology, Sylhet-3114; Department of Chemistry, Mawlana Bhashani Science and Technology University, Santosh, Tangail-1902; Bioinformatics Division, National Institute of Biotechnology, Ganakbari, Ashulia, Savar, Dhaka-1349; Department of Computer Science and Engineering, Bangladesh University of Engineering and Technology, Dhaka - 1000; Department of Genetic Engineering and Biotechnology, East West University, Dhaka-1212; Department of Microbiology, Noakhali Science and Technology University, Noakhali – 3814; Molecular Biotechnology Division, National Institute of Biotechnology, Ganakbari, Ashulia, Savar, Dhaka-1349

**Keywords:** Rotavirus A, VP4 protein, RNA interference, siRNA design, Molecular simulation

## Abstract

Rotavirus is a major cause of severe diarrheal disease in children under the age of five, with reduced vaccine effectiveness in low-resource settings causing substantial morbidity and mortality. In the absence of approved antiviral therapeutics, treatment is largely supportive, urging the need for targeted and precision-based interventions. VP4 protein plays an essential role in viral attachment, entry, and infectivity, making it a suitable target for targeted therapy. In this context, RNA interference is a specific method for inhibiting viral gene expression with its efficacy depending on sequence conservation, target accessibility, and compatibility with the RISC-loading machinery. In the present study, an integrative *in silico* approach was employed to design and evaluate siRNAs targeting conserved regions of the VP4 gene across six geographically diverse countries. Candidate siRNAs were screened using established design rules and regression-based scoring with off-target filtering. Three optimized siRNAs were further assessed through structural modeling, molecular docking, and molecular dynamics simulations to examine interactions with human Dicer, TRBP, and Argonaute-2. Comparative dynamic analyses identified one siRNA with enhanced structural compatibility, reduced conformational fluctuations, and stable interactions with RISC-loading proteins. These findings provide a rational computational basis for VP4-targeted siRNA development, facilitating experimental validation.

## Introduction

Acute rotavirus diarrheal diseases is a leading cause of mortality and morbidity among children under 5 years old in low- and middle-income countries (LMICs) ^1^. Rotavirus A is the most prominent etiological agent of infantile diarrhea, responsible for an estimated 122,000–215,000 deaths between 2013 and 2017, about 37% of all diarrheal deaths worldwide ^2,3^. Rotavirus A causes acute gastroenteritis by infecting enterocytes of the small intestine, as well as some central nervous system complications and biliary atresia ^4,5^. Immunization with live attenuated vaccines such as RotaTeq and Rotarix has reduced disease burden in high-income regions, yet vaccine effectiveness remains lower in low- and middle-income countries. Contributing factors include enteric co-infections, malnutrition, maternal antibody interference, and viral genetic heterogeneity. Moreover, no licensed antiviral therapeutics specifically targeting rotavirus are currently available, and clinical management remains largely supportive ^6–9^.

Rotavirus A, a member of the Reoviridae family, possesses an 11-segment double-stranded RNA (ds RNA) genome encoding six structural and six non-structural proteins ^10^. Among these, the outer capsid spike protein VP4 is a critical determinant of viral attachment, receptor engagement, membrane penetration, and infectivity ^11^. Proteolytic cleavage of VP4 by trypsin into VP5 and VP8 activates cell entry functions, and VP4 is a major neutralization antigen ^6^. It is involved in binding to cellular receptors, penetration, hemagglutination, virulent pathogenicity, and neutralization ^12^. Collectively, these properties render VP4 an attractive molecular target for antiviral intervention

RNA interference (RNAi) is a biological process in which RNA molecules inhibit gene expression, typically by causing the destruction of specific mRNA molecules, offering a programmable and sequence-specific approach to suppress gene expression at the post-transcriptional level ^13^. Small interfering RNAs (siRNAs), upon dicer processing and incorporation into the RNA-induced silencing complex (RISC), guide argonaute-2–mediated cleavage of complementary mRNA transcripts ^14^. Unlike small-molecule inhibitors that target viral proteins post-translation, siRNA therapeutics inhibit viral gene expression upstream at the transcript level, thereby preventing protein synthesis ^15^. RNAi-based antiviral strategies have demonstrated proof-of-concept efficacy against diverse RNA viruses including FMD virus ^16^, Human papilloma virus ^17,18^, Hepatitis B virus ^19^, Polio Virus ^20^, Influenza ^21^ and so on, highlighting their therapeutic versatility. Previous studies have demonstrated that siRNAs targeting rotaviral genes, including VP4, can reduce viral replication and progeny production ^22,23^. However, systematic computational prioritization integrating cross-strain sequence conservation, thermodynamic optimization, structural modeling, RISC-loading compatibility, and dynamic stability analysis remains underexplored. Given the high mutation rate of RNA viruses, conservation-guided design combined with structural validation is essential to enhance translational robustness.

Here, we report an integrative in silico framework to identify structurally optimized siRNAs targeting conserved regions of the VP4 gene across geographically diverse Rotavirus A strains. By combining rule-based design algorithms, regression-based scoring, mRNA accessibility prediction, molecular docking with human Dicer, TRBP, and Argonaute-2, and molecular dynamics simulations, we systematically evaluated sequence efficacy, structural compatibility, and dynamic stability within the RISC-loading machinery. This multi-tiered strategy enabled the rational prioritization of a lead siRNA candidate with favorable thermodynamic and structural attributes, providing a robust computational foundation for subsequent experimental validation and potentially transformative antiviral development.

## Results

### Prediction and Selection of siRNA

Multiple sequence alignment of VP4 gene sequences from six geographically diverse rotavirus A strains revealed conserved areas appropriate for siRNA targeting. A total of 38 siRNAs were predicted using the conserved region sequence as input in the siDirect server and were further manually analyzed to find the most suitable siRNA for silencing VP4 gene that surpassed the recommended threshold value of each of the specified algorithms. The data is provided in **Supplementary Data 1**. Finally, three siRNAs were found that are listed in **Table 1**.

**Table 1.** Chosen siRNAs comprising the sense and anti-sense strand sequences, GC content and scores obtained from rule-based methods.

| Given Name | siRNA target within mRNA | Sense 5' – 3' | Anti-sense 5'- 3' | % GC | Ui-Tei (Ia/Ib) | Amarzguioui (≥3) | Reynolds (≥6) |
| --- | --- | --- | --- | --- | --- | --- | --- |
| siRNA 01 | CGATTTAG | CGAUUUA | UUA AACU | 31.6 | Ib | 4 | 7 |
|  | G TTCAGTT | GGUUCAG | GAACCUA |  |  |  |  |
|  | TAACA | UUUA Aca | AAUCGua |  |  |  |  |
| siRNA 02 | GGTAAATG | GGUAAA U | UGUAGUU | 31.6 | Ib | 4 | 6 |
|  | ATTCAACT | GAUUCAA | GAAUCAU |  |  |  |  |
|  | ACAGT | CUACAgu | UUACCuc |  |  |  |  |
| siRNA 03 | GTATATGA | GUAUAUG | UAUUCGU | 36.8 | Ib | 3 | 6 |
|  | GAGCACGA | AGAGCAC | GCUCUCA |  |  |  |  |
|  | ATAAT | GAAUAau | UAUACua |  |  |  |  |

The table shows the sense (5' → 3') and anti-sense (5' → 3') strand sequences for three siRNAs designed to target specific mRNA sequences, along with their GC content. The table also includes efficacy scores from three rule-based methods: Ui-Tei (which is used for predicting the efficacy of siRNAs based on various parameters), Amarzguioui, and Reynolds.

### Target Accessibility and Thermodynamics Analysis

The secondary structure (**Figure 1**) prediction of VP4 gene revealed variable local accessibility across the transcript. The results showed that siRNA 1 and 3 have the highest accessibility to target mRNA suggesting favorable target site exposure (**Table 2**). The i-score was determined using the scoring matrix of a linear regression algorithm which is provided in **Supplementary Data 4**.

**Figure 1.**
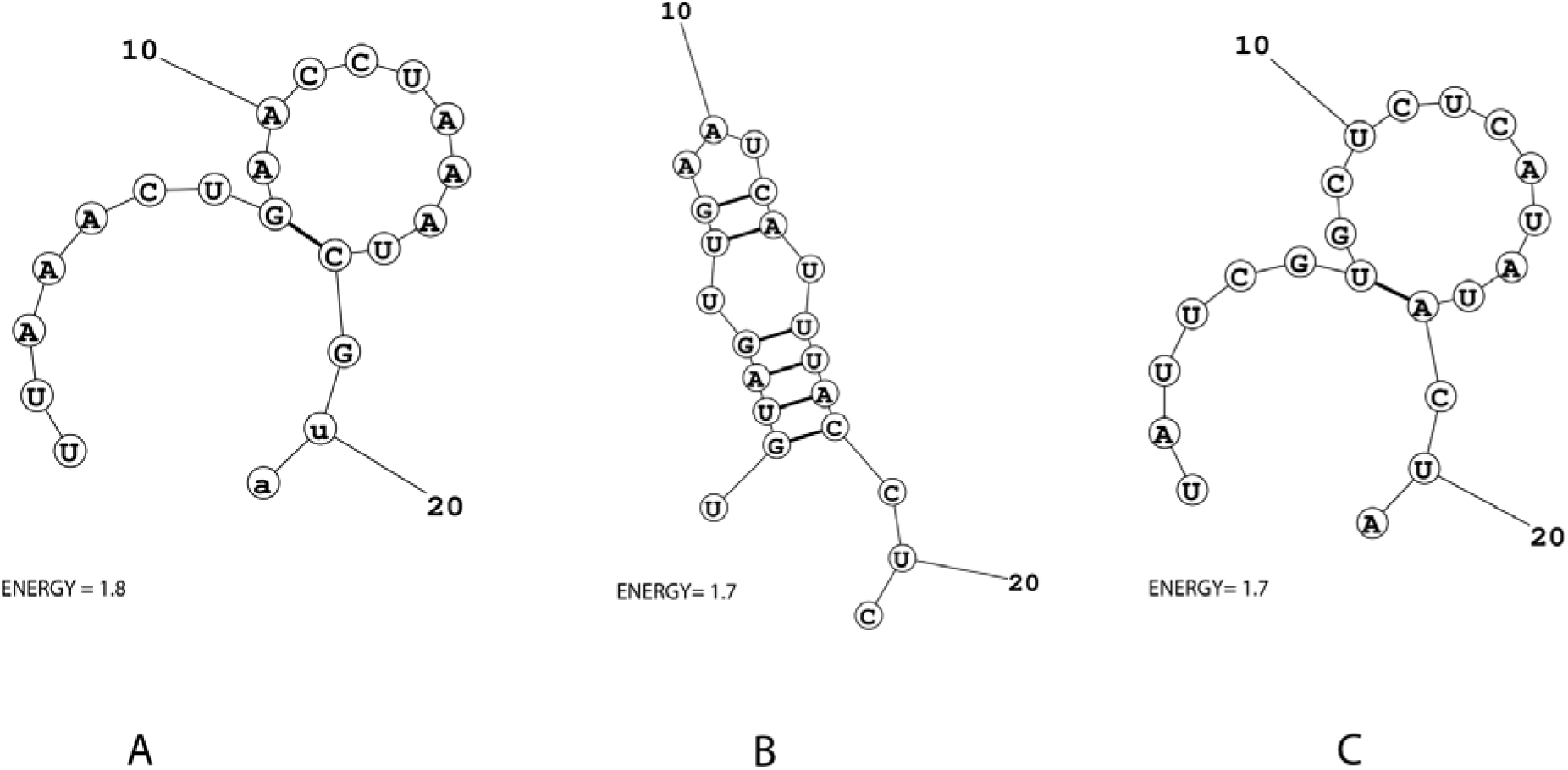
Predicted secondary conformations of the three designed siRNAs (A) siRNA01, (B) siRNA02 and (C) siRNA03. Structures were generated using the MaxExpect tool and are depicted with canonical Watson–Crick base pairing. Nucleotides are shown as labeled circles, with 5′ and 3′ termini indicated, and stem–loop features highlighting intramolecular base pairing patterns.

**Table 2.** The i-score, target accessibility score and MaxExpect calculated Energy Score of selected siRNAs.

| Given name | i-Score ($\geq 66$) | Target accessibility score ($\geq 12$) | MaxExpect calculated Energy Score for guide strands |
| --- | --- | --- | --- |
| siRNA 01 | 85.48 | 18.17 | 1.8 |
| siRNA 02 | 80.52 | 17.73 | 1.7 |
| siRNA 03 | 70.32 | 18.42 | 1.7 |

The i-score ( $\geq 66$ ), target accessibility score ( $\geq 12$ ), and MaxExpect-calculated energy score for the guide strands of three selected siRNAs are parameters used to evaluate the efficacy and stability of siRNA candidates in gene silencing experiments. The siRNA 01 exhibits the highest i-score and target accessibility score, while all three siRNAs show comparable energy scores, hinting similar efficiency in strand pairing and stability.

All the siRNAs had an internal temperature below 65°C and heterodimer binding free energies within the optimal range for effective silencing, ΔG value between -35 and -25 kcal/mol, which ensures better performance. According to our data, siRNA03 had the lowest ΔG value (-30.92 kcal/mol), indicating significant duplex formation. siRNA01 and siRNA02 had balanced thermodynamic profiles, making them ideal for RISC-mediated processing. The three siRNAs were then selected for further analysis.

### Structural Modeling of selected siRNAs

Three-dimensional models of the designed siRNAs were generated using RNAComposer, and the resulting structures were obtained in Protein Data Bank (PDB) format. They were visualized using PyMOL (**Figure 2**)

**Figure 2.**
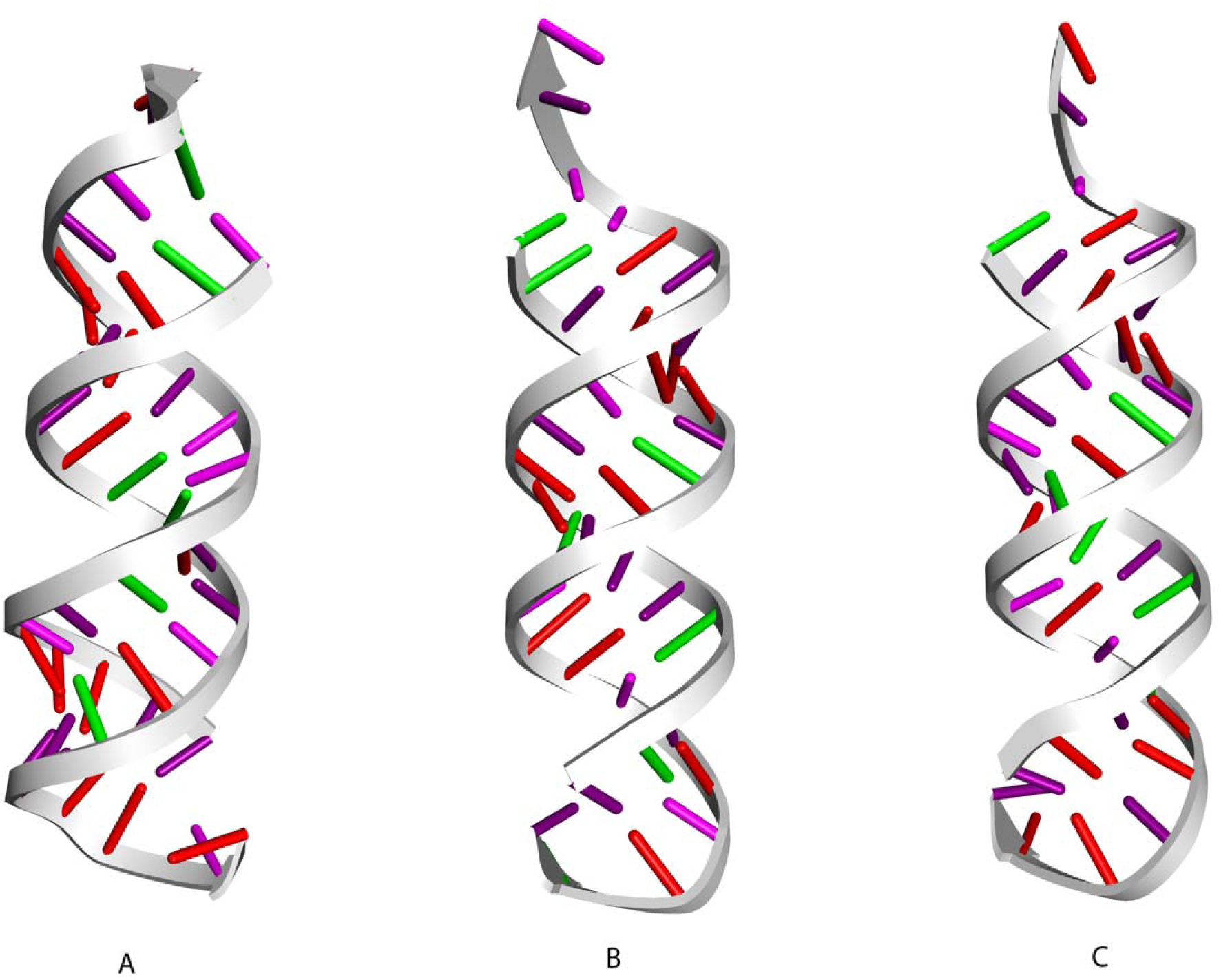
Three-dimensional structural models of the designed siRNAs showing (A) siRNA 01 (B) siRNA 02 and (C) siRNA 03. The models illustrate the canonical A-form RNA helical geometry, with the phosphate backbone shown as a continuous ribbon and individual nucleotide bases depicted as colored sticks. These structures were generated using Discovery Studio Visualizer to evaluate overall helical integrity, base stacking, and structural consistency among the siRNA candidates prior to functional assessment.

### Molecular Docking Analysis

An evaluation of the binding affinities of each of the selected siRNAs with the major components of the RISC-Loading complex was carried out showing the docking scores which were consistent across all the proteins where the control siRNAs showed the strongest interactions with the most negative binding energies, as shown in **Table 4**. The designed siRNAs showed weaker but favorable affinities.

Molecular docking of the 3 computationally designed siRNAs were performed with the human dicer along with the control. The control siRNA displayed the strongest interaction score (-383.12 kcal/mol). Among the designed candidates, siRNA01 showed comparatively higher affinity, whereas siRNA02 and siRNA03 exhibited weaker interactions. **Fig. 3** depicts the interactions between the dicer and the control siRNA, as well as the three generated siRNAs. All the protein-ligand interface data are provided in **Supplementary Data 2**.

**Figure 3.**
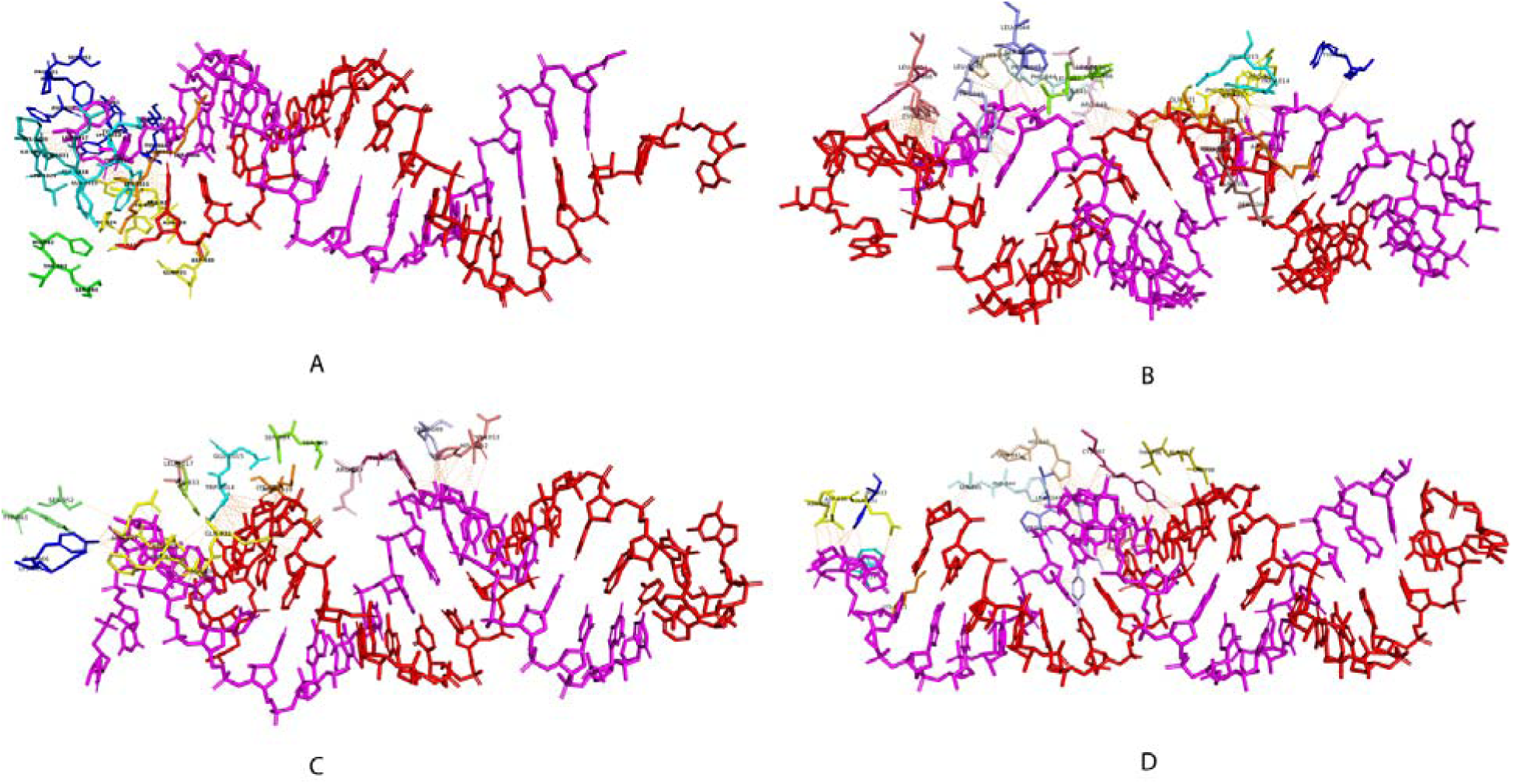
Molecular interaction profiles between human Dicer and (A) Control (B) siRNA 01 (C) siRNA 02 and (D) siRNA 03. The siRNA molecules are shown as stick representations, while interacting amino acid residues of Dicer are highlighted and labeled at the binding interface. Dashed lines indicate predicted non-covalent interactions, including hydrogen bonds and electrostatic contacts, that stabilize each complex.

Molecular docking analyses with the trans-activation response RNA-binding protein (TRBP) indicated that the reference siRNA formed the most stable complex, as reflected by the lowest binding energy. Among the designed candidates, siRNA01 exhibited the strongest affinity (– 418.99 kcal/mol), followed by siRNA03 (–411.23 kcal/mol) and siRNA02 (–404.13 kcal/mol). **Fig. 4** shows the residues of TRBP involved in interactions with the control siRNA and each of the three designed siRNAs.

**Figure 4.**
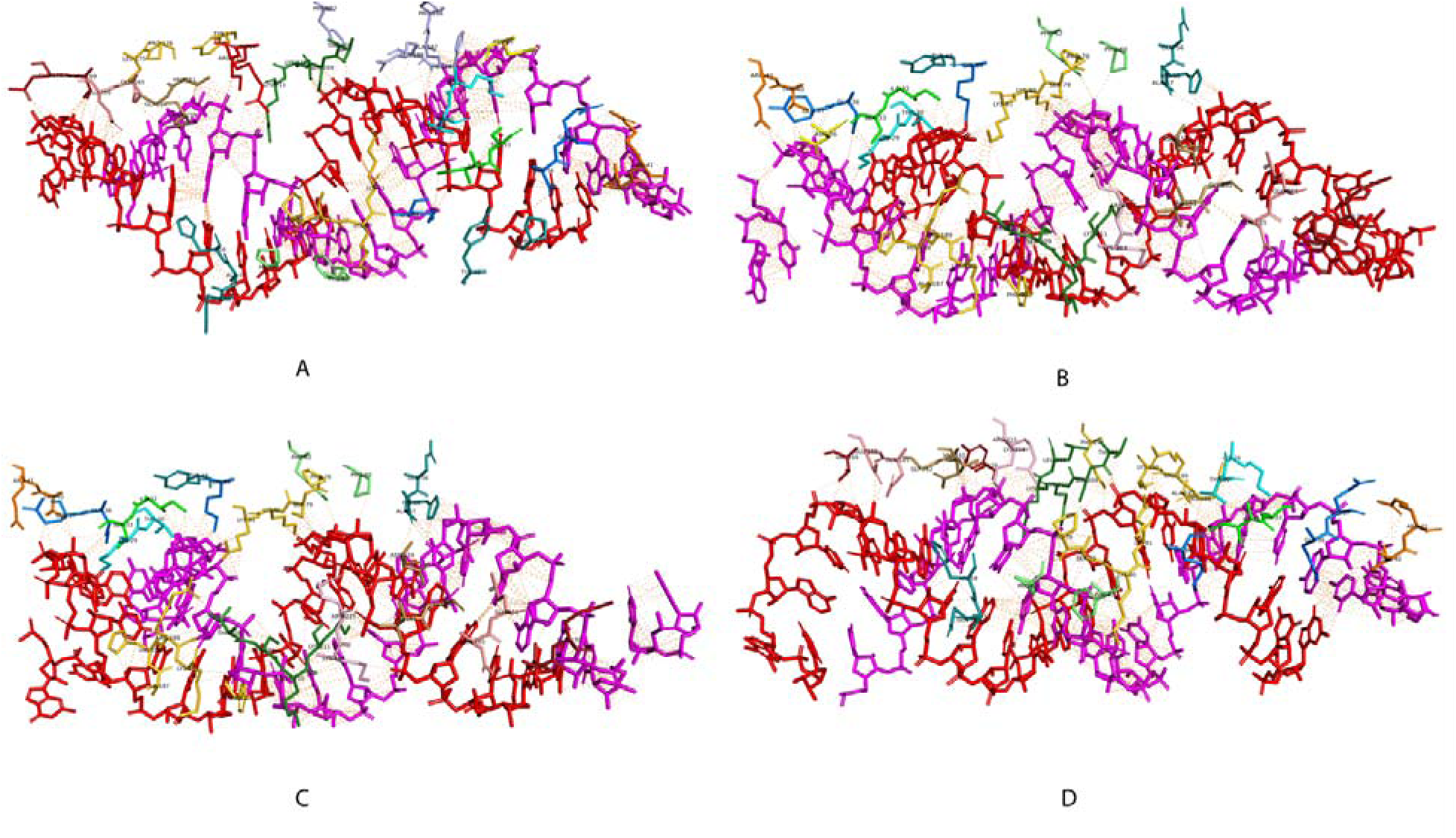
Molecular interaction profiles between TRBP and (A) Control (B) siRNA 01 (C) siRNA 02 and (D) siRNA 03. The siRNA molecules are shown as stick representations, while interacting amino acid residues of TRBP are highlighted and labeled at the binding interface. Dashed lines indicate predicted non-covalent interactions, including hydrogen bonds and electrostatic contacts, that stabilize each complex.

At last, we performed molecular docking of the designed siRNA guide strands with human Argonuate-2. The control siRNA had the most negative docking score (–300.66kcal/mol) showing similar pattern to that of human Dicer and TRBP. The designed siRNAs exhibited weaker interactions with low but favorable docking scores, with siRNAs 02 and 03 outperforming siRNA01. Minimum overlap in interaction residues between the control and tailored siRNAs was detected (**Figure 5**).

**Figure 5.**
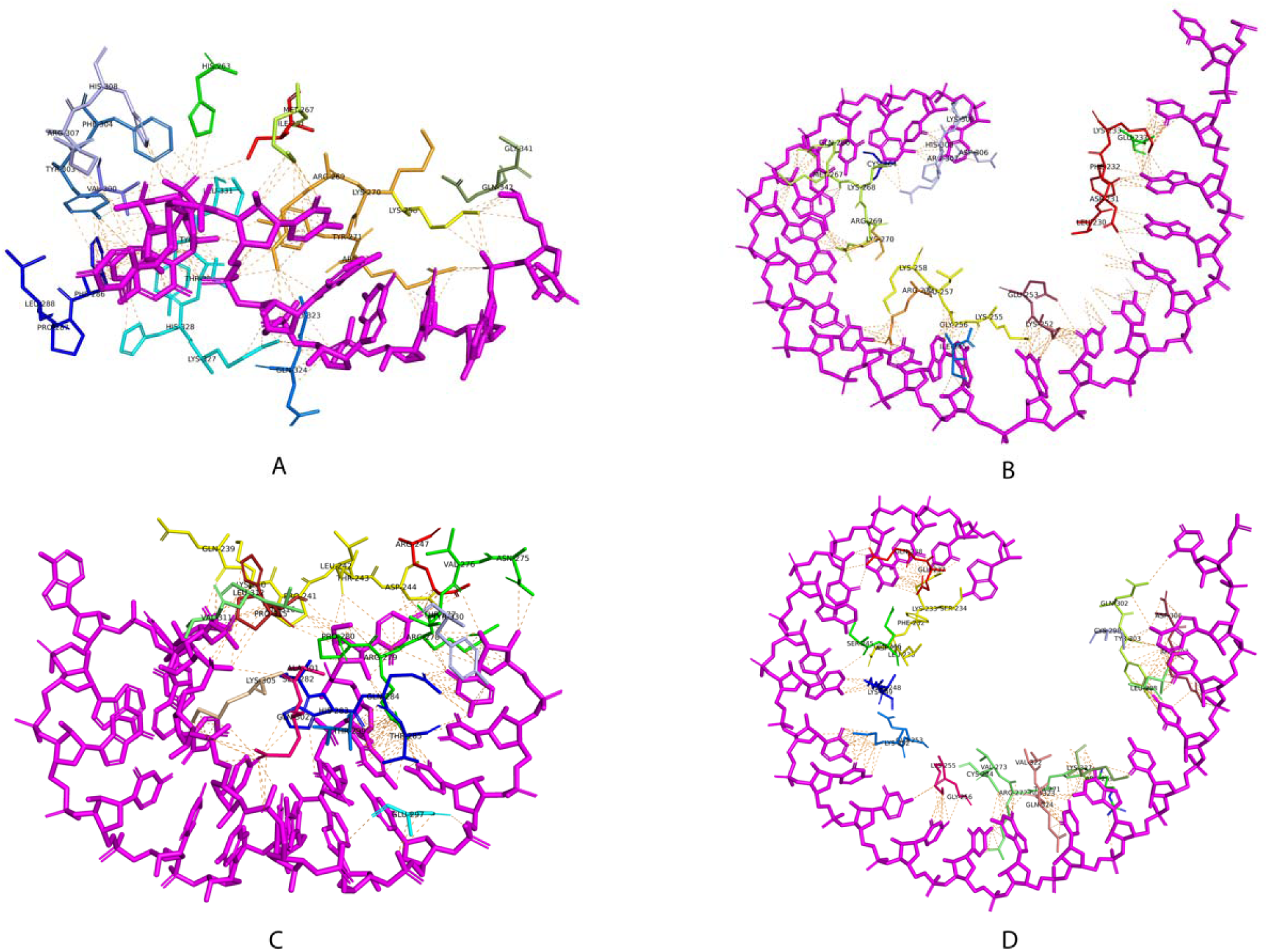
Molecular interaction profiles between Argonuate-2 and guide strands of (A) Control (B) siRNA 01 (C) siRNA 02 and (D) siRNA 03. The siRNA molecules are shown a stick representations, while interacting amino acid residues of TRBP are highlighted and labeled at the binding interface. Dashed lines indicate predicted non-covalent interactions, including hydrogen bonds and electrostatic contacts, that stabilize each complex.

### Molecular Dynamics Simulation (MDS) results

Molecular dynamics simulations were carried out for the designed siRNA–protein complexes, together with relevant controls, in order to examine their behavior at an atomic level and assess complex stability under conditions simulating the physiological environment. For Dicer, the siRNA01 complex consistently exhibited reduced SASA (around 147 to 155 nm^2^) and Rg (around 2.05-2.10 nm) values compared to the control and other siRNA complexes, indicating a more compact and stable interaction (**Figure 6**). RMSD analysis showed that both the control and siRNA01 complexes rapidly reached equilibrium and maintained low deviations of around 0.25 – 0.32 nm, whereas siRNA02 displayed moderate instability and siRNA03 showed pronounced structural fluctuations. RMSF profiles further confirmed reduced residue-level flexibility at the Dicer binding interface for siRNA01, while siRNA03 induced elevated fluctuations in key loop regions. Comparative dynamic metrics consistently favored siRNA01, which exhibited reduced conformational fluctuation and enhanced complex compactness relative to the other candidates.

**Figure 6.**
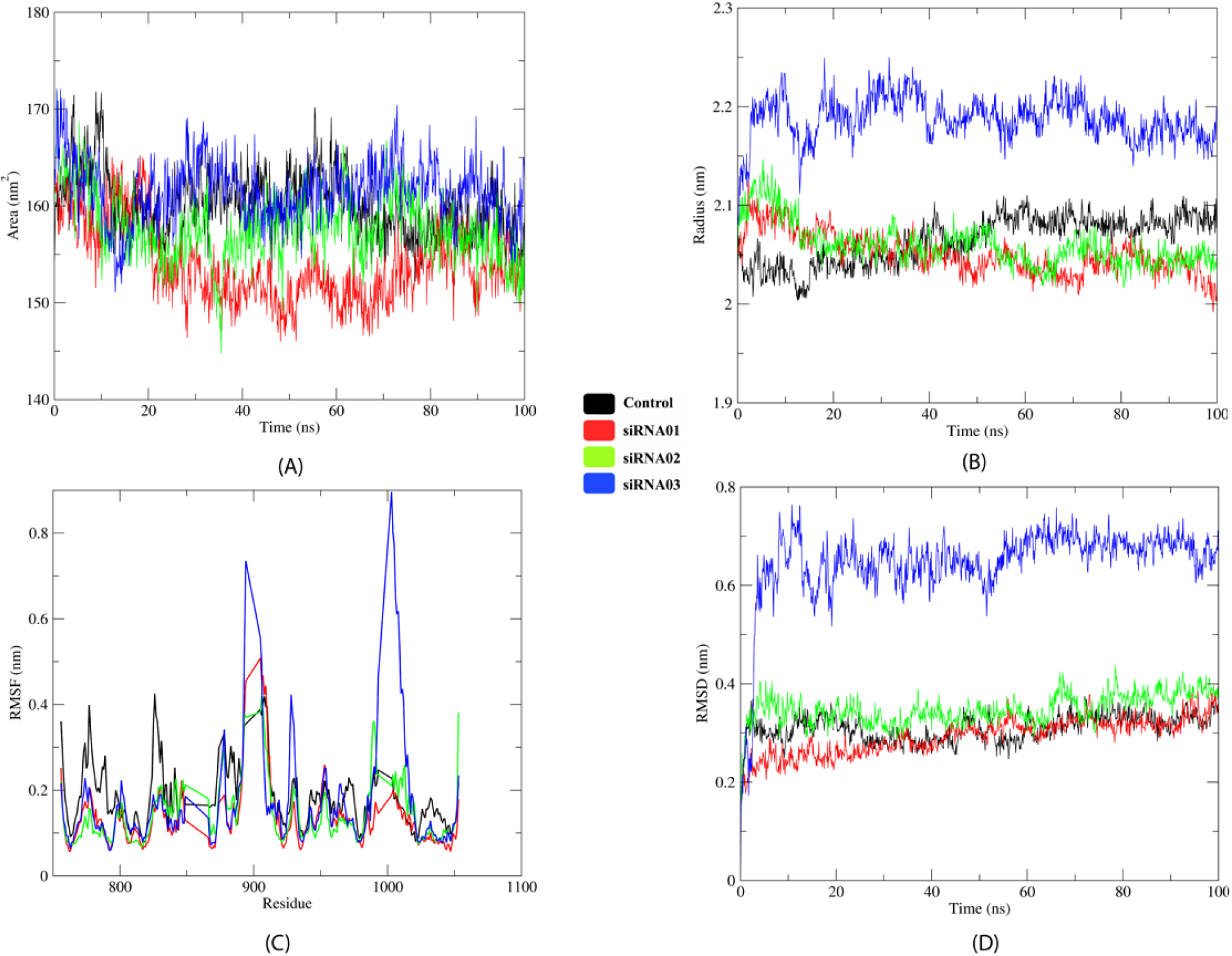
Molecular dynamics simulation outcomes for Dicer-siRNA complexes. Comparative analyses of human Dicer in complex with control siRNA and the designed siRNA candidates (siRNA01, siRNA02, and siRNA03) over a 100 ns simulation period showing A) Solvent-accessible surface area (SASA), (B) radius of gyration (Rg), (C) residue-wise root mean square fluctuation (RMSF), and (D) root mean square deviation (RMSD) over a 100-ns simulation, illustrating differences in temporal variations in protein surface exposure upon siRNA binding, global compactness, local flexibility and structural stability among th complexes.

In TRBP complexes, all complexes retained their overall fold showing only differences based on the siRNA they formed a complex with (**Figure 7**). However, siRNA03 consistently increased solvent exposure, structural expansion, and local flexibility relative to the control. siRNA01 and siRNA02 demonstrated moderate stability, with siRNA01 maintaining comparatively lower SASA and Rg values and reduced residue fluctuations.

**Figure 7.**
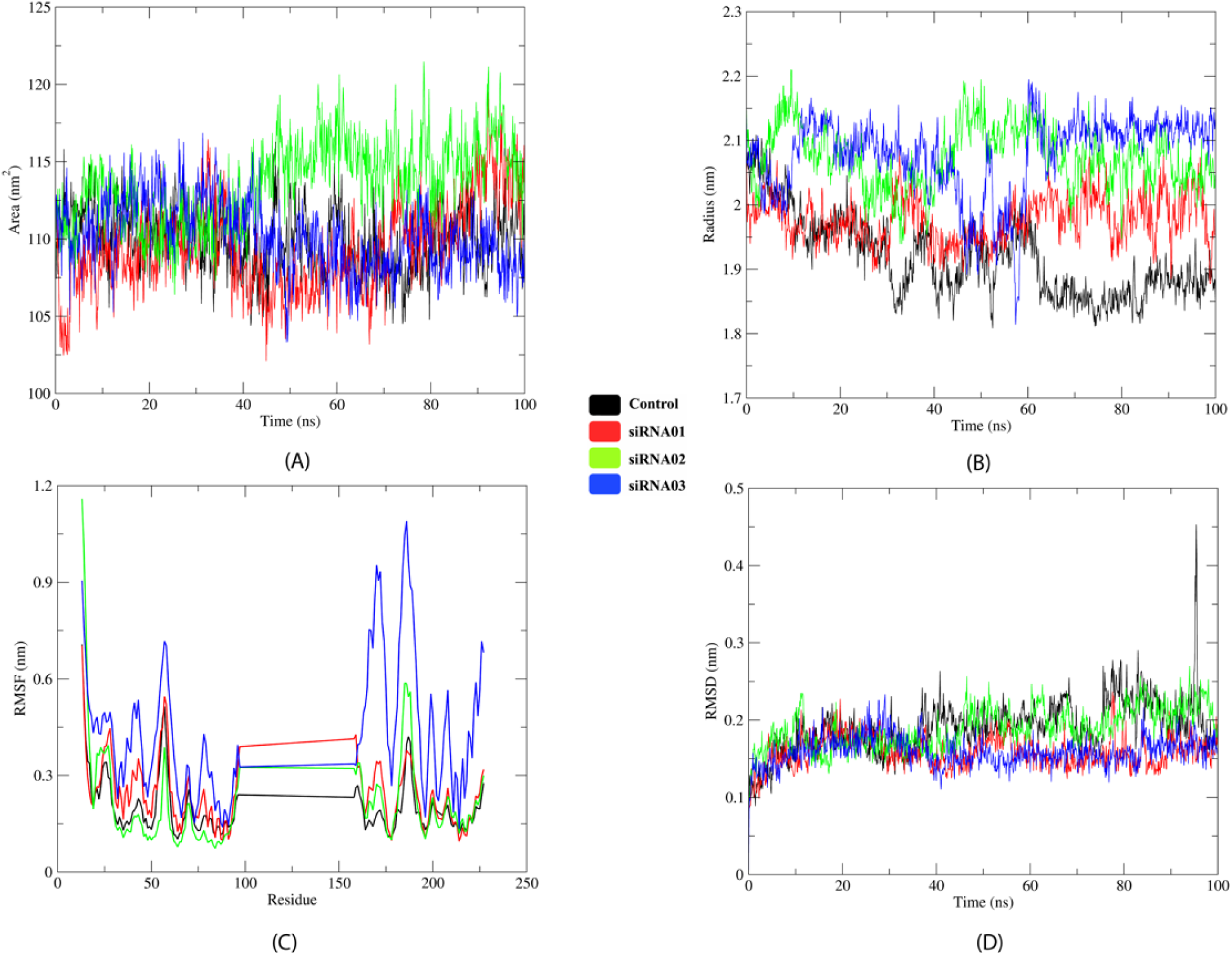
Molecular dynamics simulation outcomes for TRBP-siRNA complexes. Comparative analyses of TRBP in complex with control siRNA and the designed siRNA candidates (siRNA01, siRNA02, and siRNA03) over a 100 ns simulation period showing A) Solvent-accessible surface area (SASA), (B) radius of gyration (Rg), (C) residue-wise root mean square fluctuation (RMSF), and (D) root mean square deviation (RMSD) over a 100 ns simulation, illustrating differences in temporal variations in protein surface exposure upon siRNA binding, global compactness, local flexibility and structural stability among th complexes.

On the contrary, Ago2 proved to be highly robust towards all the docked siRNAs **(Figure 8)**. RMSD and RMSF values indicated preservation of the global fold, with only minor differences among siRNA complexes. siRNA01 showed slightly reduced solvent exposure and conformational variability, whereas siRNA02 exhibited marginally increased adjustments. Overall, all the siRNAs displayed proper accommodation with the Ago2 with siRNA01 having consistent stabilizing advantage over the others.

**Figure 8.**
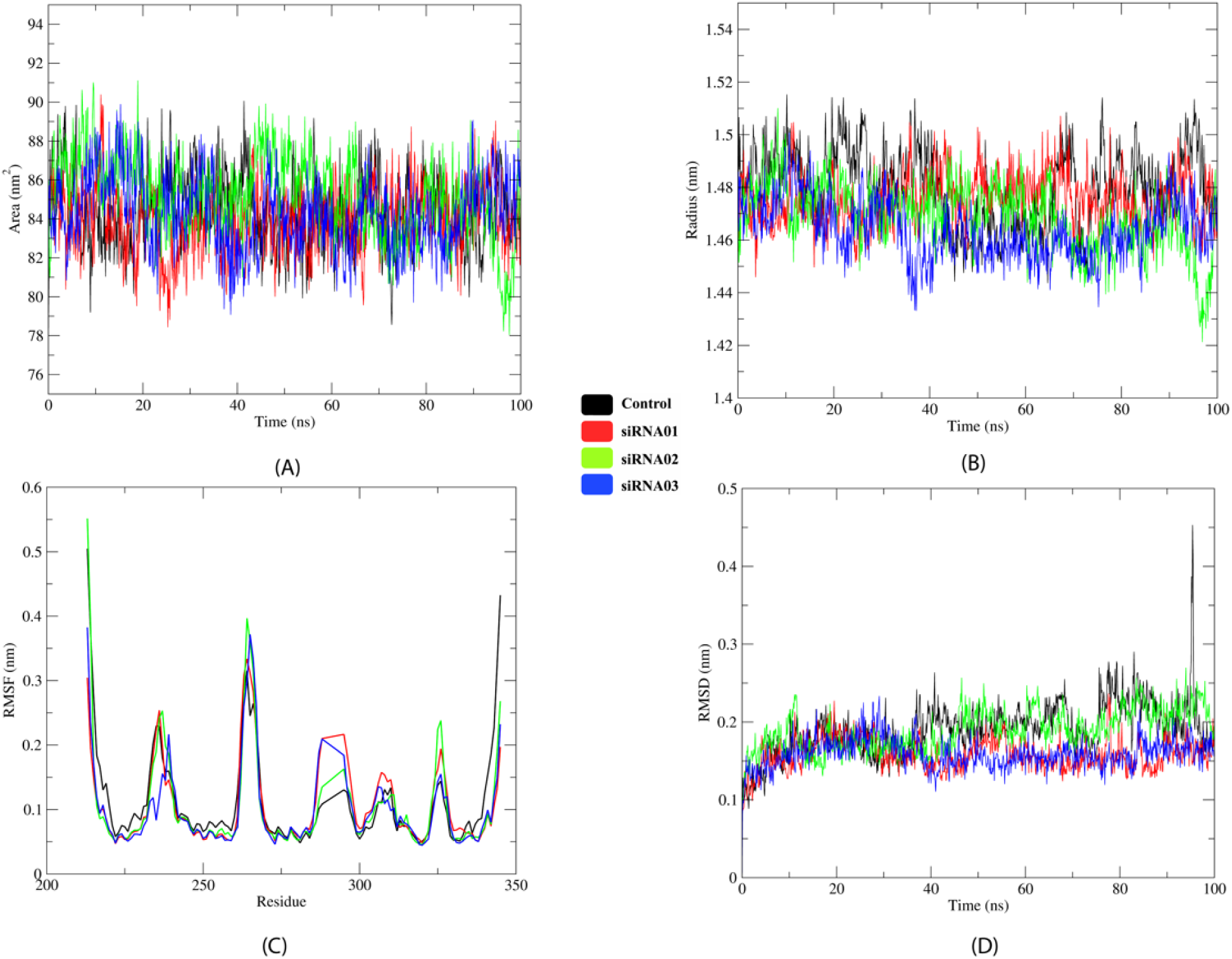
Molecular dynamics simulation outcomes for Ago2-siRNA complexes. Comparative analyses of TRBP in complex with control siRNA and the designed siRNA candidates (siRNA01, siRNA02, and siRNA03) over a 100 ns simulation period showing, A) Solvent-accessible surface area (SASA), (B) radius of gyration (Rg), (C) Residue-wise root mean square fluctuation (RMSF), and (D) Root mean square deviation (RMSD) over a 100 ns simulation, illustrating differences in temporal variations in protein surface exposure upon siRNA binding, global compactness, local flexibility and structural stability among th complexes.

Analysis of the RMSD, Rg, local flexibility and solvent exposure across all the complexes revealed siRNA01 to repeatedly appear as the most stable and structurally compatible siRNA, while siRNA02 showed acceptable but weaker behavior in comparison. The siRNA03 showed consistency in disrupting the protein dynamics revealing suboptimal binding. The analysis supports siRNA01 being the most promising candidate for processing by the human Dicer while having stronger and consistent TRBP-siRNA01 interaction ensuring a stable loading into Ago2 which corresponds to the MID and PAZ domains that anchor the 5’ and 3’ ends of the guide strand required for effective gene silencing.

## Discussion

The persistent burden of rotavirus-associated gastroenteritis in developing countries, despite vaccine deployment, necessitates complementary antiviral strategies capable of overcoming vaccine performance disparities ^2^. In this context, small interfering RNAs (siRNAs) represent a promising therapeutic strategy, as they can selectively suppress the expression of disease-associated genes through sequence-dependent mechanisms ^26^.

In the present study, we employed a conservation-driven computational pipeline to design siRNAs targeting the VP4 gene across strains originating from geographically distinct regions. Inclusion of sequences from South Asia, East Asia, and multiple African regions includes, Bangladesh, Wuhan in China, India, Malawi in East Africa, Togo in West Africa, Gauteng province in South Africa minimized strain bias and increased the probability of broad-spectrum efficacy. We performed multiple sequence alignment (MSA) analysis of the sequences to determine the conserved region, which was used for designing three siRNAs. Conservation-guided targeting is critical for RNA viruses, where rapid mutation can compromise sequence-specific interventions.

The selected siRNAs were then evaluated using widely used rule based and machine learning filters (Ui-Tei rules, Amarzguioui rules, Reynolds rules, and i-Score), which is consistent with prior studies that targeted viral genes and oncogenes ^25^. Maintenance of GC content within the optimal 30–60% range balanced duplex stability with efficient strand unwinding during RISC loading ^26^. Target accessibility analysis incorporated mRNA secondary structure considerations, recognizing that sterically occluded regions may impair guide strand hybridization and cleavage efficiency. Among the candidates, siRNA01 consistently achieved superior composite scores, indicating favorable predicted silencing efficiency (**Figure 1**).

The candidates were further analyzed based on their thermodynamic properties which further supported the functional potential of the designed siRNAs. All selected siRNAs exhibited internal melting temperatures below 65 °C and heterodimer binding free energy (ΔG) values that fell within the favorable range of −25 to −35 kcal/mol **(Table 3)**. These thermodynamic characteristics reflect the energetic landscape surrounding nucleic acid interactions and are relevant to polycationic binding as well as intracellular release behavior ^36,37^. The availability of the target location inside the mRNA molecule as well as the potentiality of a chosen siRNA candidate determine how well it silences genes. Therefore, a key criteria for determining the effectiveness of siRNA-mediated gene silencing is siRNA target site accessibility ^38^.The siRNA01 and siRNA03 showed a target accessibility score of above 18 **(Table 2)**.

**Table 3.** Thermodynamic properties of the designed siRNAs.

| Given Name | Internal melting temperature, $T_m$ ($^{\circ}\text{C}$) | $\Delta G$ for heterodimer binding (kcal/mol) |
| --- | --- | --- |
| siRNA 01 | 46.8 | -26.66 kcal/mol |
| siRNA 02 | 48.1 | -27.81 kcal /mol. |
| siRNA 03 | 51 | -30.92 kcal/mol |

Summary of important thermodynamic parameters for the three designed siRNAs, including internal melting temperature ( $T_m$ , $^{\circ}\text{C}$ ) and Gibbs free energy ( $\Delta G$ , kcal · mol $^{-1}$ ) associated with sense and antisense heterodimer formation. The melting temperature depicts duplex thermal stability, whereas the $\Delta G$ values indicate favorable energy of siRNA strand pairing. These parameters were used to compare relative stability and binding efficiency among siRNA01, siRNA02, and siRNA03 during *in silico* analysis.

Beyond sequence and thermodynamic considerations, structural compatibility with the RNA-induced silencing complex is essential for siRNA functionality ^39^. In Molecular Docking analysis of a complex, a low and negative score usually denotes that the complex is more stable ^40^. That means that an extremely negative score would indicate a strong binding while moderately negative or a positive score corresponds to a weaker or a non-existent binding. 3D structures of the three siRNAs were modeled using RNAComposer tool which revealed appropriate hairpin free conformations minimizing undesirable self-folding tendencies. Molecular docking of these siRNAs was performed with the 3 components of the RISC-loading complex, human Dicer, TRBP and argonuate-2 proteins. The control siRNAs present in these structures were isolated and separately docked with the proteins to compare and validate the docking results of the selected siRNAs. In most cases, the docking results were remarkably close to the control **(Table 4).** The siRNA 01 showed the most promising result with all the complexes. A number of common interactions were also found between the siRNA and all the proteins **(Figure 3 – 5).**

**Table 4.** Docking scores obtained for interactions between designed siRNAs and the proteins of the RISC-Loading complex.

| Complex | Docking scores (kcal/mol) |
| --- | --- |
| Dicer-control | -383.12 |
| Dicer-siRNA01 | -306.24 |
| Dicer_siRNA02 | -279.67 |
| Dicer_siRNA03 | -280.75 |
| TRBP Control-siRNA | -480.72 |
| TRBP-siRNA01 | -418.99 |
| TRBP-siRNA02 | -404.13 |
| TRBP-siRNA03 | -411.23 |
| Ago2-control | -300.66 |
| Ago2-siRNA01 | -243.57 |
| Ago2-siRNA02 | -252.43 |
| Ago2-siRNA03 | -276.86 |

Docking-derived binding energies (kcal/mol) for complexes formed between the designed siRNA candidates (siRNA01, siRNA02, and siRNA03) and key components of the RNA-induced silencing complex (RISC) complex, including Dicer, TRBP, and Ago2. Corresponding protein-control siRNA interactions are shown for comparison with more negative docking scores indicating stronger predicted binding affinity, providing a comparative assessment of siRNA compatibility with RISC proteins during *in silico* interaction analysis.

MD simulation, on the other hand assessed the stability of the siRNAs with the docked protein complexes. It provided a deeper insight into the structural dynamics of the complexes to evaluate conformational stability under near-physiological conditions ^41^. As such, key parameters indicating the structural stability and molecule behavior was analyzed **(Figure 6**–**8).** The radius of gyration showed the compactness of our siRNAs while the Root Mean Square Deviation (RMSD) showed the stability as they were found to show consistent across the protein complexes. SASA, on the other hand, is the surface area of a molecule accessible to the solvent. A lower SASA value is an indication that the surface is less open implying that the molecule is more compact. The RMSF measures the fluctuation of the atoms or residues of the complexes with a high RMSF indicating more flexibility. ^42^ The siRNA 01 had the most consistent attributes across all the parameters. In Dicer complexes, siRNA01 reduced local flexibility and maintained low RMSD values, hinting efficient accommodation within the PAZ and platform domains essential for siRNA processing. TRBP complexes showed moderate tolerance to all siRNAs, although siRNA03 induced increased local fluctuations, indicating subpar stability. In contrast, Ago2 displayed remarkable robustness across all complexes, with siRNA01 again exhibiting marginally enhanced compactness and reduced solvent exposure, supporting efficient guide strand anchoring within the MID and PAZ domains. Overall, the implementation of MDS in our studies added a depth and more insight to the dynamics of our siRNAs, simulating a biological environment. A comprehensive understanding of its dynamic behavior gave more credibility to our developed siRNAs for gene silencing of the VP4 protein.

Despite the strengths of this integrative computational approach, several limitations must be acknowledged. *In silico* predictions cannot fully model intracellular pharmacokinetics, endosomal escape efficiency, nuclease degradation, innate immune activation (e.g., Toll-like receptor engagement), or potential viral escape mutations. Additionally, docking-derived energy values represent relative scoring metrics rather than experimentally validated binding free energies. Therefore, experimental validation in rotavirus-infected cell culture systems— incorporating quantitative viral load assessment, protein expression analysis, and cytotoxicity profiling—will be essential to confirm antiviral efficacy.

Nevertheless, this study establishes a comprehensive computational workflow integrating conservation analysis, thermodynamic profiling, structural docking, and molecular dynamics simulations to rationally prioritize siRNA candidates targeting rotaviral VP4. The identification of siRNA01 as a dynamically stable and RISC-compatible candidate provides a mechanistically supported lead for downstream experimental validation and potential therapeutic development. More broadly, this framework may be adapted for rapid siRNA design against other emerging RNA viral pathogens.

## Conclusion

In this study, we establish a conservation-guided computational framework for the rational design and structural prioritization of siRNAs targeting the VP4 gene of Rotavirus A. Integration of sequence optimization, thermodynamic profiling, molecular docking, and molecular dynamics simulations identified siRNA01 as a structurally stable and RISC-compatible lead candidate.

The convergence of conservation, binding affinity, and dynamic stability metrics supports its potential for efficient guide strand incorporation and functional silencing. While experimental validation remains necessary, this integrative in silico strategy significantly refines candidate selection and accelerates the translational pathway for RNAi-based antivirals targeting rotavirus.

## Material and methods

The overall workflow is shown in **Figure 9**

**Figure 9.**
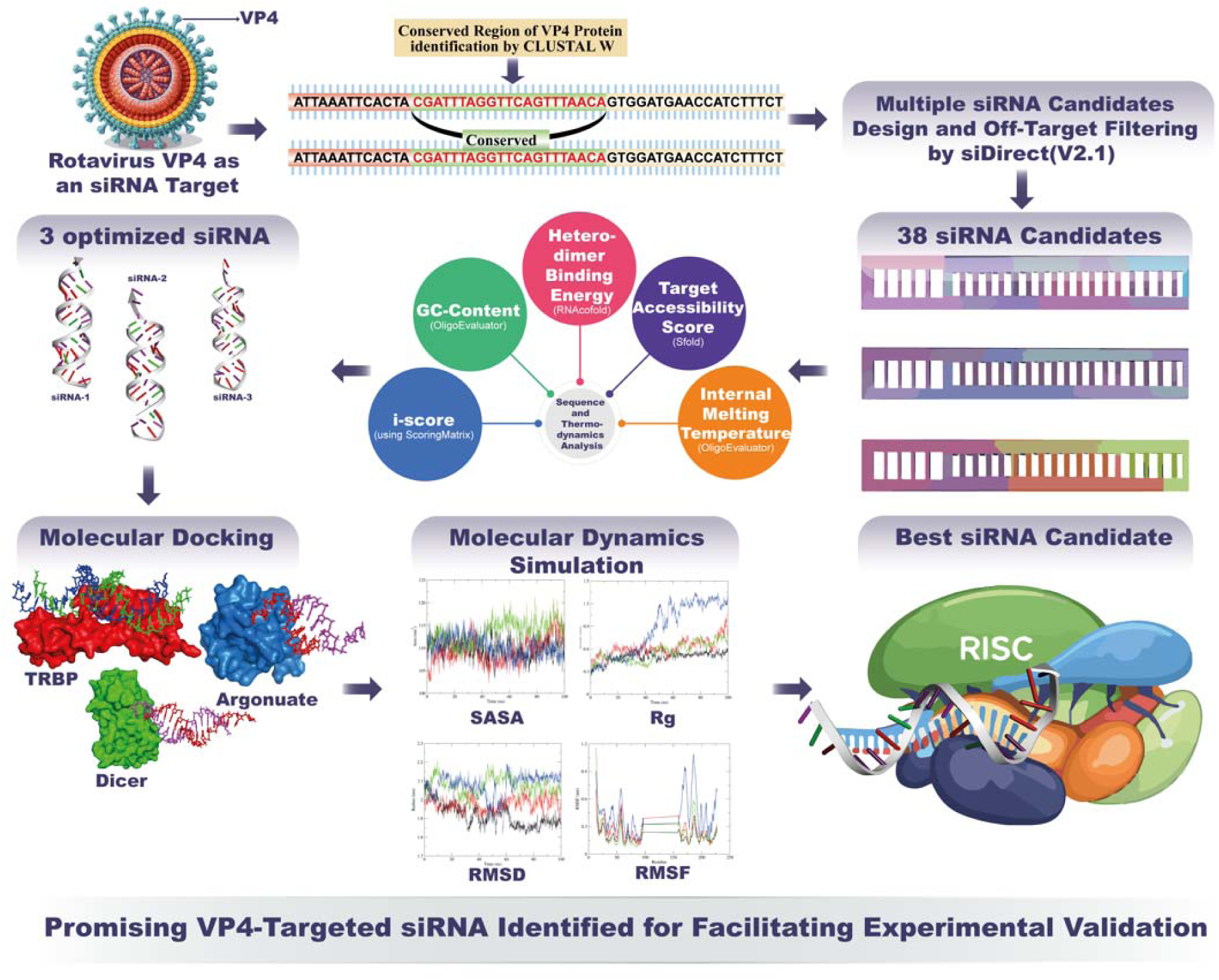
An overall workflow. The schematic outlines of the process of identifying optimized siRNAs targeting the conserved region of rotavirus VP4. Using siDirect (V2.1), 38 siRNA candidates are designed and filtered for off-target effects. Three optimized candidates are selected based on factors such as GC-content, dimer binding energy, and target accessibility. The best candidate is then validated through molecular docking with Dicer, TRBP and Argonaute-2, followed by molecular dynamics simulations to assess stability.

### Sequence Analysis and siRNA Design

VP4 coding sequences representing six geographically (Bangladesh, Wuhan in China, India, Malawi in East Africa, Togo in West Africa, Gauteng province in South Africa) distinct strains were retrieved from NCBI (https://www.ncbi.nlm.nih.gov/). Multiple sequence alignment was done using ClustalW ^24^ to find the conserved region amongst all the sequences.

The siDirect version 2.1 web server was used to design potential siRNA for the study ^25^ applying Ui-Tei, Amarzguioui, Reynolds, and Tuschl design rules. For seed-duplex stability, a Tm (melting temperature) of 21.5 degree Celsius was set as the maximum and the G/C content was set between 30 % to 60% ^26^. The i-score (inhibitory score) was calculated using scoring parameters through a simple algorithm which was previously used to detect siRNAs applying a linear regression model to 2431 siRNAs ^27^. The scoring matrix is provided on **Supplementary Data 4**. Off-target homology was assessed using BLAST against the human transcriptome and non-target viral sequences against the entire GenBank database (https://blast.ncbi.nlm.nih.gov/Blast.cgi) ^28^, and siRNAs with more than 15 consecutive nucleotide matches to non-target sequences were removed.

### Structural and Thermodynamic Characterization of Designed siRNAs

The G/C content of the developed siRNA and the internal melting temperature (Tm) for the sense strand of each potential siRNA duplex were analyzed using the OligoEvaluator analysis tool (http://www.oligoevaluator.com) ^29^. The secondary structure was predicted with the MaxExpect tool (https://rna.urmc.rochester.edu/RNAstructureWeb/Servers/MaxExpect/MaxExpect.html) ^30^. The Sfold web server’s siRNA module (http://sfold.wadsworth.org) was used to calculate the target accessibility score of the selected siRNAs ^31^. Thermodynamic stability of siRNA –mRNA heteroduplexes was evaluated utilizing the RNAcofold web server (http://rna.tbi.univie.ac.at/cgi-bin/RNAWebSuite/RNAcofold.cgi) ^32^.

### Structural Modeling and Molecular Docking

The RNAfold webserver was used to generate secondary structures of the siRNAs in dot-bracket notation (Vienna format) for three-dimensional modeling with RNAComposer through the motif-template based method (http://rnacomposer.ibch.poznan.pl/Home). The PDB file was downloaded and the 3D structure was visualized using PyMOL ^33^.

Molecular docking analyses were performed to evaluate interactions between the designed siRNAs and key components of the RISC-loading complex, including transactivation response element RNA-binding protein (TRBP), Dicer, and Argonaute-2. The corresponding crystal structures were obtained from the Protein Data Bank (RCSB) ^34^. The crystal structures that were collected for docking included the human Dicer Platform–PAZ–Connector Helix cassette bound to a 17-mer siRNA with 5′-phosphate and UU-3′ termini (PDB ID: 4NGF; 3.1 Å resolution), TRBP dsRBD domains 1 and 2 complexed with a 19-bp siRNA (PDB ID: 5N8L), and the PAZ domain of human Argonaute-2 (residues 214–347) in complex with the RNA sequence CGUGACUCU (PDB ID: 6RA4). All structures were retrieved from the RCSB Protein Data Bank. The binding residues of the human dicer were determined using the control siRNA and docked our designed siRNAs based on those residues. The docking analyses were conducted using the HDOCK server.

### Molecular dynamics (MD) simulation

MD simulations for the designed siRNAs, siRNA-protein complexes and relevant controls were done by employing CHARMM36 force field formed by the CHARMM-GUI server which was run with the GROMACS 2018.3 program ^35^. A periodic cubic simulation box was generated where the siRNA was placed within a 1.0 nm distance between the complex and edges of the solvated box. The systems were then neutralized by adding counter ions. Energy minimization was carried out using the steepest descent method, followed by an equilibration phase of 100 ps under constant volume and temperature (NVT) conditions. System equilibration was done using NPT ensemble for 1 ns in the subsequent phase. An identical simulation protocol was applied for the simulation of siRNAs in their end state for about 1 ns. All the simulation used a time step of 2 fs and was run at 300K temperature and one atmospheric pressure, which replicated standard experimental conditions. Subsequently, standard GROMACS functions evaluated the Solvent Accessible Surface Area (SASA), Radius of gyration (Rg), Root Mean Square Fluctuation (RMSF), and Root Mean Square deviation (RMSD). All simulations and analyses were executed on the high-performance computing cluster of the Bioinformatics Division at the National Institute of Biotechnology, Bangladesh.

## Acknowledgements

We are grateful to Bioinformatics Division and its computational facilities for extending this research work.

## Author contributions

Abu Nayeem Ahmed and Khaleda Jahan Satu: Writing – original draft, Visualization, Software, Methodology, Formal analysis, Data curation, Conceptualization. A.B.Z. Naimur Rahman, SM Sajid Hasan, Mohammad Nazmus Sakib, Md. Ekram Hossan: Writing–original draft, Methodology, Formal analysis, Conceptualization. Arittra Bhattacharjee and Zeshan Mahmud Chowdhury: Software, Validation, Review & editing. Ziaul Faruque Joy: Data curation, Visualization. Md. Jahirul Islam: Review & editing. Mohammad Uzzal Hossain: Supervision, Conceptualization, Review & editing.

## Data availability

All data generated or analyzed during this study are included in this published study.

